# Gap-free nuclear and mitochondrial genomes of *Ustilaginoidea virens* strain JS60-2, a fungal pathogen causing rice false smut

**DOI:** 10.1101/2022.10.25.513643

**Authors:** Yin Wang, Lei Yang, Qun Yang, Jie Dong, Yufu Wang, Yuhang Duan, Weixiao Yin, Lu Zheng, Wenxian Sun, Jing Fan, Chao-Xi Luo, Guotian Li

**Affiliations:** State Key Laboratory of Agricultural Microbiology, Hubei Hongshan Laboratory, the Provincial Key Laboratory of Plant Pathology of Hubei Province, College of Plant Science & Technology, Huazhong Agricultural University, Wuhan 430070, China; Key Lab of Horticultural Plant Biology, Ministry of Education, College of Plant Science and Technology, Huazhong Agricultural University, Wuhan 430070, China; College of Plant Protection and the Ministry of Agriculture Key Laboratory of Pest Monitoring and Green Management, China Agricultural University, Beijing, 100193, China; State Key Laboratory of Crop Gene Exploration and Utilization in Southwest China, Sichuan Agricultural University, Chengdu, Sichuan 611130, China

**Keywords:** *Villosiclava virens*, *Oryza sativa*, Nanopore, mycotoxins, whole genome sequencing

## Abstract

Rice false smut (RFS), caused by *Ustilaginoidea virens*, has become a major disease in recent years, and mycotoxins produced by *U. virens* often threaten food safety. To study fungal pathogenesis and identify potential targets for developing new fungicides, gap-free nuclear and complete mitochondrial genomes of *U. virens* JS60-2 were sequenced and assembled. Using the second and third generation sequencing data, we assembled a 38.02-Mb genome that consists of seven contigs with the contig N50 being 6.32-Mb. In total, 8,486 protein-coding genes were annotated in the genome, including 21 secondary metabolism gene clusters. We also assembled the complete mitochondrial genome, which is 102,498 bp, with 28% GC content. The JS60-2 genomes assembled in this study will facilitate research on *U. virens* and contribute to RFS control.

## Genome Announcement

Rice false smut (RFS), caused by the filamentous fungus *Ustilaginoidea virens*, has become one of the most devastating diseases in major rice-growing areas (Sun et al., 2020). RFS causes up to 49% of yield losses, when the environmental conditions are favorable (Neelam et al., 2022). Additionally, multiple types of mycotoxins are produced by *U. virens*, such as ustiloxins and ustilaginoidins, that are toxic to plants, animals and humans (Li et al., 2019). *U. virens* initially infects rice floral organs, and, in the later infection phase, blackish green false smut balls develop in rice spikelets (Zhang et al., 2014). Chlamydospores and sclerotia produced by *U. virens* can survive on rice stubbles and paddy field weeds for months. When the temperature and humidity are suitable in the new growing season, chlamydospores germinate and produce conidia that are important for primary infection. Genetic resistance is the most effective strategy in disease control, and some quantitative trait locis (QTLs) against RFS have been identified, such as loci *qFsr10* and *qFsr12* (Zhou et al., 2014). However, effective resistance genes for RFS have not been cloned (Sun et al., 2020), and chemical control is still one important approach for RFS control. To study fungal pathogenesis and further identify potential targets for developing new fungicides, it is indispensable to assemble a high-quality *U. virens* genome. Therefore, we sequenced and assembled gap-free nuclear and mitochondrial genomes of *U. virens* strain JS60-2.

*U. virens* strain JS60-2 was isolated from Jiangsu Province, China. We chose *U. virens* JS60-2 for genome sequencing because it is a local *U. virens* strain that is suitable for screening local rice varieties for resistance and it can also infect Kitaake, Nipponbare, Wanxian98, Zhonghua11, and other rice varieties that are commonly used in research. We extracted DNA from JS60-2 mycelia and amplified the internal transcribed spacer (ITS) region (accession number MH938066.1). Compared with sequences in the National Center for Biotechnology Information (NCBI) database, the JS60-2 ITS region is identical to 41 *U. virens* strains. The second- and third-generation sequencing technologies (Wang et al., 2021) were used for genome sequencing of the JS60-2 strain, and they were performed on the PromethION platform and the BGISEQ-500 platform, respectively. We obtained 3.8 Gb of short reads and 7.3 Gb of long reads through the second and third technologies, and we then used NextDenovo (available on GitHub) to assemble the JS60-2 genome. Subsequently, the draft genome was polished using short and long reads (Wang et al., 2022). Finally, a 38.02-Mb gap-free JS60-2 genome was assembled(Table 1), which consists of seven contigs (N_50_, 6.32 Mb). These seven contigs show high collinearity with reference genomes UV-8b and UV-P1 (Supplementary Figs. S1 and S2). Each contig matches one of seven chromosomes using D-genes (Cabanettes and Klopp, 2018). Using RAGOO (Alonge et al., 2019), these contigs were anchored into seven chromosomes, and telomere sequences in chromosomes 2 and 4 were identified for both chromosomal ends. Furthermore, repetitive sequences were found and masked, using RepeatModeler (v2.0.1) (Flynn et al., 2020) and Repeatmasker (v4.0.7) (Tarailo-Graovac and Chen, 2009). BUSCO (v4.1.4) (Seppey et al., 2019) was used to analyze the completeness of the genome assembly, and results showed that 98.80% of genes were represented, indicating a high-quality assembly of the JS60-2 genome. The JS60-2 genome is the first gap-free *U. virens* genome, and the contig N_50_ has been improved in comparison to previous *U. virens* genomes (Zhang et al., 2021; Pramesh et al., 2020; Kumagai et al., 2016).

**Table 1.** Summarized genomic information of the *U. virens* strain JS60-2

| Variables | Statistics |
| --- | --- |
| Sequence coverage | 216 X |
| Genome size (Mb) | 38.02 |
| Contigs | 7 |
| Contig N50 size (bp) | 6,315,893 |
| Max contig size (bp) | 7,452,382 |
| Number of predicted genes | 8,468 |
| Secreted proteins | 649 |
| Predicted effectors | 222 |
| BUSCO completeness (%) | 98.80 |

We obtained 10.5 Gb of RNA-seq data from mycelia cultured under light condition, mycelia cultured under dark condition and chlamydospore balls on inoculated rice spikelets. (Zhao et al., 2022). In total, 8,486 protein-coding genes were predicted, and 7,202 (84.87%) of them were annotated, using eggNOG (Huerta-Cepas et al., 2019). Additionally, 649 putative secreted proteins were identified by an established bioinformatic pipeline (Dong et al., 2015), 222 of which are candidate effectors, including 101 predicted apoplastic effectors and 121 cytoplasmic effectors. AntiSMASH (Blin et al., 2021) was used to predict the secondary metabolism gene clusters. The JS60-2 genome contains five polyketide synthases (PKS), four nonribosomal peptide synthetases (NRPS), five NRPS-like fragment (NPRS-Like), two non-alpha poly-amino acids synthetase, and five terpene synthases. Among them, one PKS biosynthesis gene cluster located in chromosome 1 was predicted to synthesize an ACR toxin and one NRPS biosynthesis gene cluster located in chromosome 4 for a nonribosomal peptide. The former may contribute to fungal infection (Izumi et al., 2012), and the latter may confer *U. virens* tolerance to reactive oxygen species (Chen et al., 2013).

The whole mitochondrial genome that facilitates monitoring the population genomic structure of the pathogenwas assembled using the second-generation sequencing short reads (Fig. 1). The size of the JS60-2 mitochondrial genome is 102 kb, and the GC content is 28%. We used BlastN (Chen et al., 2015) to predict two types of ribosomal RNAs including *rns* and *rnl. Rps3*, an important ribosome-related gene maintaining normal mitochondrial functions, was also predicted in the *rnl* intron region. BlastX was used to predict 14 proteins that are involved in oxidative phosphorylation and producing ATPs. Using tRNAscan-SE_v2, 29 tRNAs were predicted (Chan et al., 2021). In contrast to *U. virens* UV-8b, the mitochondrial genome of JS60-2 is 4.4-kb larger but the numbers of protein-encoding genes is the same (Zhang et al., 2021).

**Fig. 1.**
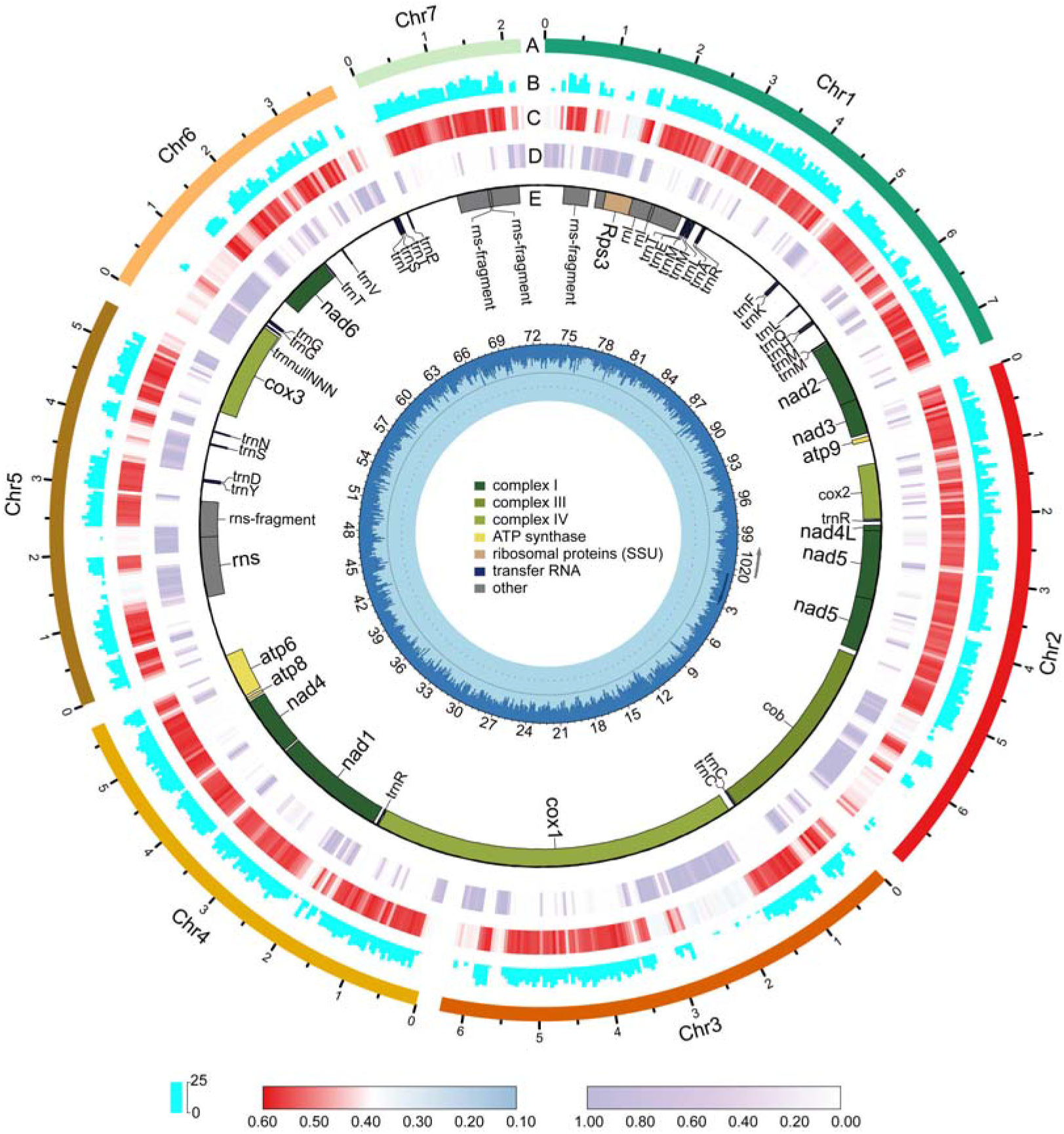
Nuclear and mitochondrial genomes of the *U. virens* strain JS60-2. (A) The seven JS60-2 chromosomes on a Mb scale. (B) Gene density. (C) GC content. (D) Repeat density. Data in nuclear genome circles are displayed in nonoverlapping 50-kb intervals. (E) Mitochondrial genome and predicted genes related to oxidative phosphorylation (green), ATP synthases (yellow), ribosomal proteins (brown), transfer RNAs (blue), and others (grey). The innermost circle shows the mitochondrial genome in the kb scale as well as the GC content.

In this study, we present a gap-free nuclear genome and a complete mitochondrial genome of *U. virens* strain JS60-2. A total of 222 effectors and 21 secondary metabolism gene clusters that may facilitate *U. virens* infection were predicted. The *U. virens* strain JS60-2 and its genomes will facilitate studies of the *U. virens-rice* pathosystem and contribute to RFS control.

## Data availability

The JS60-2 genome file is available at NCBI under the project accession number PRJNA859262, and the nuclear genome annotation file, the sequence file of candidate effectors, the mitochondrial genome file, and the mitochondrial genome annotation file have been deposited to Figshare database. All other sequencing data are available at China’s National Genomics Date Center Genome Sequence Archive (GSA) under the accession number CRA007582.

The authors declare no conflict of interest.

