## Supplemental Figure S1 and S2 for "Gap-free nuclear and mitochondrial genomes of *Ustilaginoidea virens* strain JS60-2, a fungal pathogen causing rice false smut"

### Supplemental Figures

**Figure S1** Dot plot comparing the JS60-2 and UV-8b genomes.

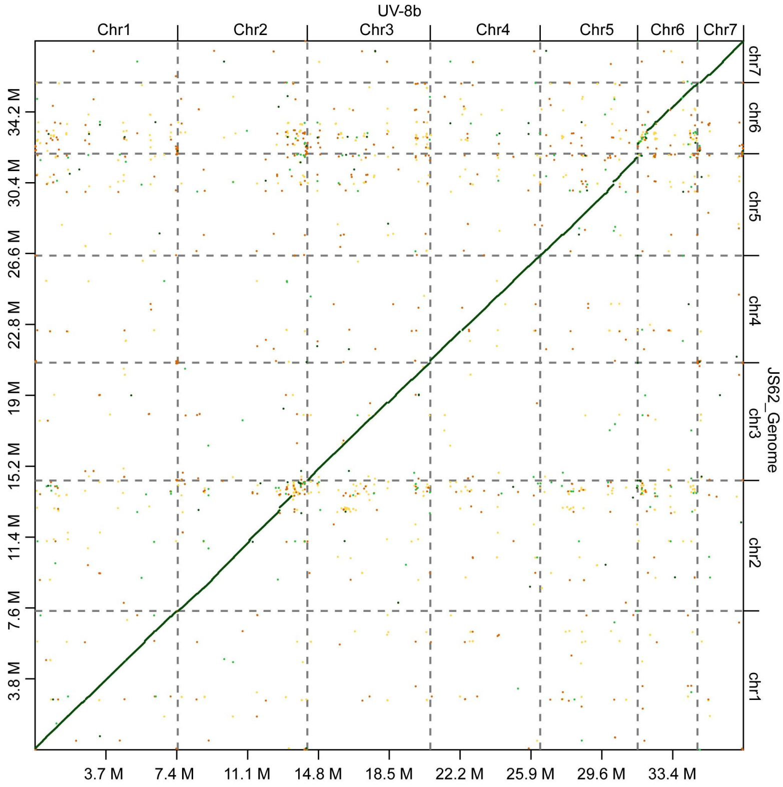


**Figure S2** Dot plot comparing the JS60-2 and UV-P1 genomes.

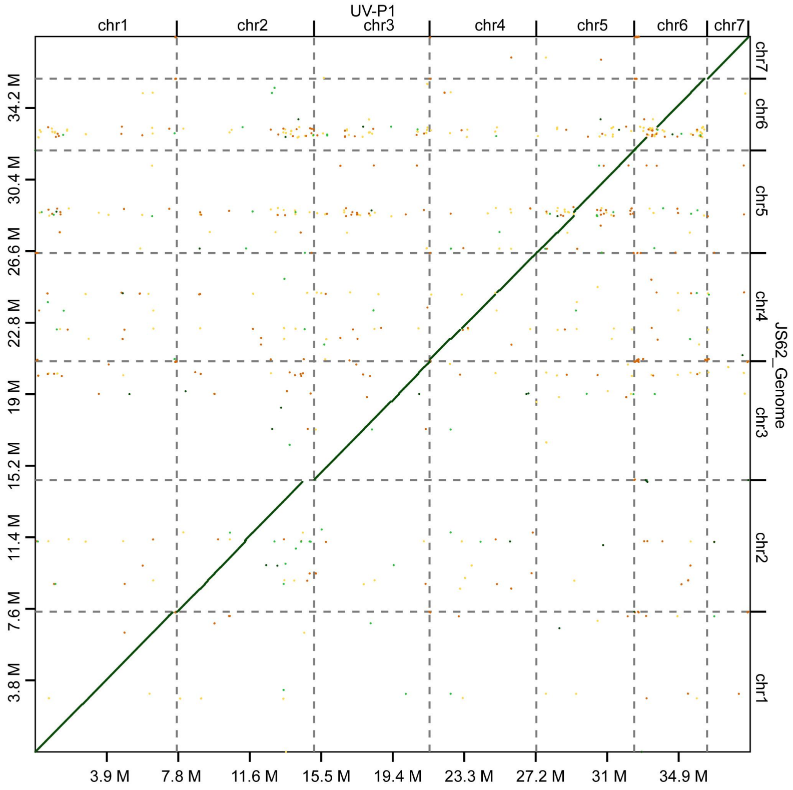
